# Endothelial Zmiz1 modulates physiological and pathophysiological angiogenesis during retinal development

**DOI:** 10.1101/2024.06.30.601426

**Authors:** Nehal R Patel, Rajan K C, Mark Y Chiang, Stryder M Meadows

**Affiliations:** Department of Cell and Molecular Biology, Tulane University, New Orleans, LA, United States; Division of Hematology-Oncology, Department of Internal Medicine, Medical School, University of Michigan, Ann Arbor, MI, United States; Tulane Brain Institute, Tulane University, New Orleans, LA, United States

**Keywords:** Zmiz1, transcription, co-factor, angiogenesis, retinopathy

## Abstract

Angiogenesis is a highly coordinated process involving the control of various endothelial cell behaviors. Mechanisms for transcription factor involvement in the regulation of endothelial cell dynamics and angiogenesis have become better understood, however much remains unknown, especially the role of non-DNA binding transcriptional cofactors. Here, we show that Zmiz1, a transcription cofactor, is enriched in the endothelium and critical for embryonic vascular development, postnatal retinal angiogenesis, and pathological angiogenesis in oxygen induced retinopathy (OIR). In mice, endothelial cell-specific deletion of *Zmiz1* during embryogenesis led to lethality due to abnormal angiogenesis and vascular defects. Inducible endothelial cell-specific ablation of *Zmiz1* postnatally resulted in impaired retinal vascular outgrowth, decreased vascular density, and increased vessel regression. In addition, angiogenic sprouting in the superficial and deep layers of the retina was markedly reduced. Correspondingly, vascular sprouting in fibrin bead assays was significantly reduced in the absence of Zmiz1, while further *in vitro* and *in vivo* evidence also suggested deficits in EC migration. In agreement with the defective sprouting angiogenesis phenotype, gene expression analysis of isolated retinal endothelial cells revealed downregulation of tip-cell enriched genes upon inactivation of *Zmiz1*. Lastly, our study suggested that endothelial Zmiz1 is critical for intraretinal revascularization following hypoxia exposure in the OIR model. Taken together, these findings begin to define the previously unspecified role of endothelial Zmiz1 in physiological and pathological angiogenesis.

## Introduction

Vasculogenesis and angiogenesis are key processes required for the initial formation and subsequent expansion of pre-existing blood vessels, respectively. Development of a functional vasculature is critical for tissue growth, regeneration, and homeostasis, while impaired vascularization is associated with various disease processes, including stroke, vascular malformations, retinopathy, and cancer^1–4^. The angiogenic generation of new blood vessels is a multi-step regulatory process intimately involving endothelial cells (ECs). For instance, vascular sprouting requires specialized ECs called tip cells that respond to both chemo-attractant and repulsive extracellular signals to guide the growth of the vasculature. Functional behaviors of ECs, such as these, are regulated by transcription factors (TFs) and transcription cofactors that mediate cell-specific gene expression changes during different phases of blood vessel growth^5,6^. However, the transcription cofactors that modulate TF involvement in the EC transcriptional program have largely remained elusive.

Zmiz1 is a transcription cofactor belonging to the protein inhibitor of activated STAT (PIAS) protein family^7^. As a transcriptional cofactor, Zmiz1 does not bind DNA directly, but instead interacts with DNA-binding TFs. Zmiz1 has been shown to regulate the transcriptional activity of multiple TFs, including P53, Androgen receptor, Smad3/4 and Notch1^8–10^. Similar to the diversity of its co-partners, Zmiz1 has been shown to have roles in various developmental processes and diseases. Zmiz1 is associated with Notch-dependent T-cell development and leukemogenesis^11^ and numerous studies point to Zmiz1 involvement in erythropoiesis^12^, osteosarcoma^13^, diabetes^14^, multiple sclerosis^15^ and in a range of neurodevelopmental disorders^16–18^. In terms of vascular development, global deletion of Zmiz1 in mice leads to lethality at embryonic day (E) 10.5 due to cardiovascular defects. Overall, the embryos are severely underdeveloped and display abnormal blood vessel development^19^. Further, microarray analysis of VEGF-deficient ECs displayed downregulation of *Zmiz1* in the gene ontology cluster associated with blood vessel development^20^ suggesting a functional role in the developing endothelium. Related, our group recently showed that Zmiz1 transcriptionally regulates lymphatic EC gene expression^21^. However, the exact function of Zmiz1 in blood vessel development and angiogenesis remains unclear.

Utilizing the murine retina, a well-established model for studying the different processes of angiogenesis^22,23^, we investigated the role of Zmiz1 in both physiological and pathological angiogenesis. In the current study, we report that loss of endothelial *Zmiz1* during embryogenesis was lethal due to vascular defects. Additionally, we demonstrated that Zmiz1 is a fundamental regulator of postnatal retinal vascular growth, including a specific and crucial function in sprouting angiogenesis. We further provided evidence of Zmiz1 involvement in the regulation of tip-cell gene expression during the vascular sprouting process. Lastly, in an oxygen induced retinopathy (OIR) mouse model, Zmiz1 was implicated in pathological neovascularization.

## Results

### Zmiz1 is expressed in the endothelium of various tissues

Given the unexplored, yet potential role of Zmiz1 in vascular development and function, we first surveyed mouse EC databases to evaluate *Zmiz1* expression in various tissues. Analysis of organ-specific EC transcriptomic data at P7 revealed highest expression of *Zmiz1* mRNA in brain ECs and comparable expression between kidney, liver, and lung ECs^24^ (Supplementary Fig. 1A). Furthermore, evaluation of single cell transcriptomic data generated using cells isolated from an adult mouse brain via fluorescence-activated cell sorting (FACS) showed a mean expression of 2.28 for *Zmiz1* in ECs^25^ (Supplementary Fig. 1B). Analysis of another single-cell transcriptomic data set using isolated brain ECs from P7 mice allowed for assessment of *Zmiz1* in different EC subtypes (arterial, venous, and capillary ECs)^24^. *Zmiz1* expression was relatively uniform across the different EC cell types (Supplementary Fig. 1C). Bulk RNA sequencing of murine retinal ECs at different developmental stages showed the presence of *Zmiz1* transcripts during early postnatal development, with highest expression levels between postnatal days (P) 6-15, and a decrease from P21-P50^26^ (Supplementary Fig. 1D). In addition, utilizing immunofluorescence analysis, we demonstrated that ZMIZ1 protein was highly enriched throughout the blood vessels of the developing P7 retina, including the arteries, veins, and capillaries (Supplementary Fig. 1E). Collectively, these studies established the presence of Zmiz1 in the endothelial lineage of postnatal and adult mouse tissues. Further, examination of human EC databases has shown *ZMIZ1* expression in both vascular and lymphatic endothelial cells^27^.

### Early endothelial cell-specific ablation of *Zmiz1* results in vascular defects and embryonic lethality

Previous work indicated that *Zmiz1* null mice die embryonically due to defects in angiogenesis^19^. To further investigate the endothelial role for Zmiz1 in embryonic vascular development, conditional Zmiz1 flox mice (*Zmiz1*^fl/fl^) were crossed to mice expressing Cre recombinase under the vascular Tie2-promoter (*Tie2*-Cre) (Fig. 1A), thereby achieving constitutive *Zmiz1*-EC deletion during embryogenesis. Homozygous deletion of Zmiz1 resulted in embryonic lethality as no viable *Zmiz1*^fl/fl^;*Tie2*-Cre (referred to as *Zmiz1*^cECKO^; constitutive EC-knockout) pups were born (Fig. 1B). Conversely, the other control genotyped pups (*Zmiz1*^fl/+^, *Zmiz1*^fl/fl^ and *Zmiz1*^fl/fl^;*Tie2*-Cre) were born at normal mendelian ratios. In timed mating, E12.5 *Zmiz1*^cECKO^ embryos displayed growth retardation and were significantly reduced in size as compared to the littermate *Zmiz1*^fl/fl^ controls (Fig. 1C, D). Between E12.5 and E13.5, mutant embryos appeared pale with a lack of blood flow in the embryo, displayed an absence of blood-filled yolk sac vessels, and/or exhibited localized hemorrhages (Fig. 1C). By E14.5 all the mutant embryos were necrotic and being reabsorbed (data not shown). PECAM-1 antibody staining of blood vessels in *Zmiz1*^cECKO^ embryos at E12.5 revealed defects in the vascular patterning of the cranial and trunk vasculature (Fig. 1E). In *Zmiz1*^cECKO^ embryos, the cranial blood vessels were not well defined and failed to form larger caliber vessels, while the dorsal vasculature, including intersomitic vessels, in the trunk were truncated and disorganized. Analysis of yolk sacs at E12.5 revealed the lack of large and distinctive blood vessels in *Zmiz1*^cECKO^ embryos in comparison to littermate controls (Fig. 1C, F). Taken together, these data showed that Zmiz1 is critical for embryonic vascular development and further indicated a major contributing role of defective angiogenesis as a cause for embryonic lethality in *Zmiz1*^-/-^ mice.

**Figure 1.**
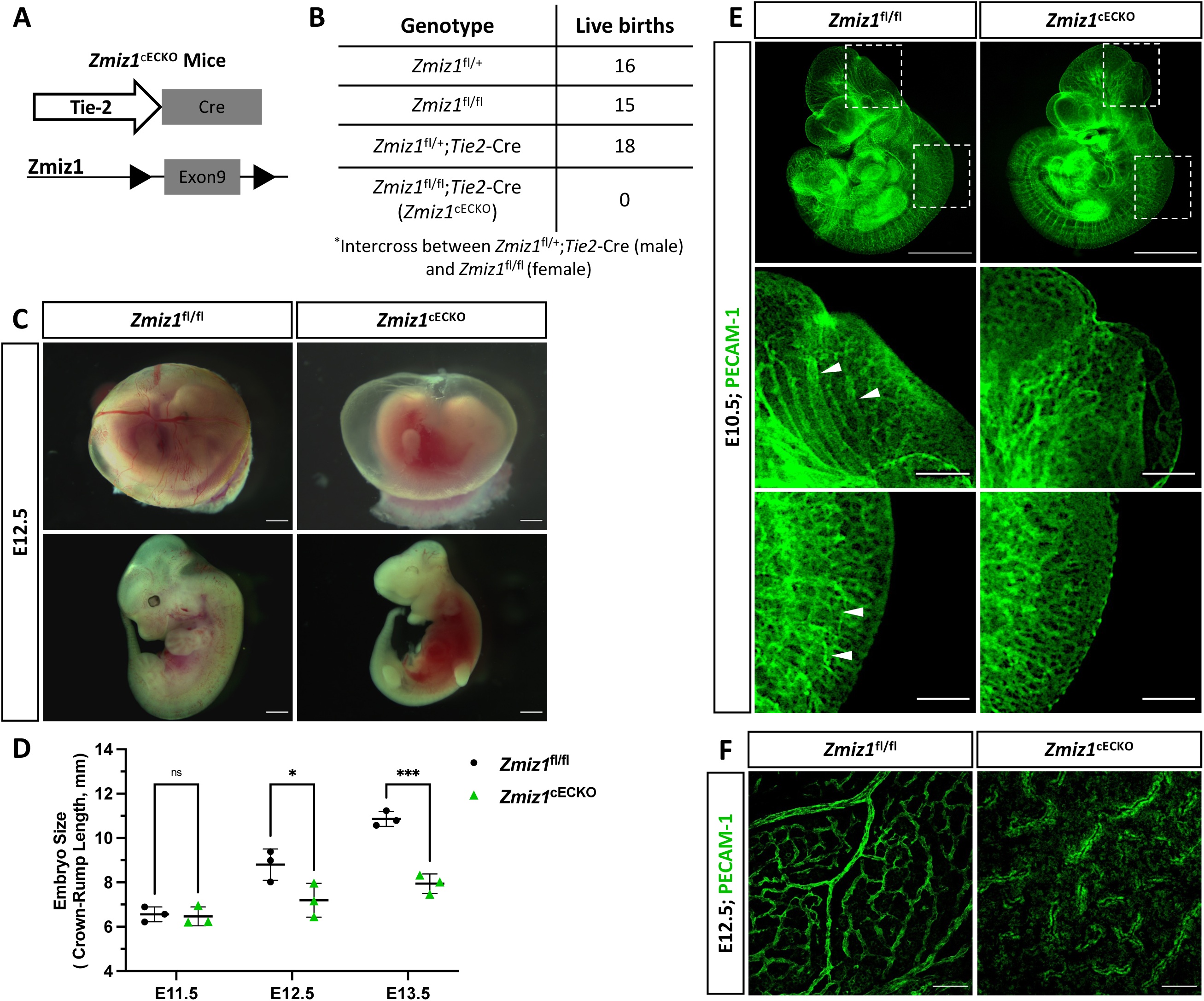
Endothelial Zmiz1 is essential for angiogenesis during embryonic development. **A,** Strategy for *Zmiz1* exon 9 deletion specifically in the ECs during embryogenesis using the Tie2-Cre mouse model (Zmiz1 constitutive endothelial cell knockout; Zmiz1cECKO). **B,** Genotypic analysis of the progeny generated by crossing *Zmiz1*^fl/+^;*Tie2*-Cre males with *Zmiz1*^fl/fl^ females at postnatal day (P) 21. All genotypes, except *Zmiz1*^fl/fl^;*Tie2*-Cre, were observed at normal Mendelian ratios. **C,** Brightfield images showing gross morphology of the intact yolk sacs and whole embryos at embryonic day (E) 12.5. Scale bars: 1 mm. **D,** Quantification of embryo size as indicated by crown-rump length from E11.5-E13.5. **E,** Whole-mount PECAM-1 immunofluorescent staining of E11.5 *Zmiz1*^fl/fl^ and *Zmiz1*^cECKO^ embryos. Scale bars 1 mm. Higher magnification of embryos shows vessel detail in the cranial and tail region (white arrowheads indicate well-organized and connected vessels in cranial and tail region). Scale bars: 100 μm. **F,** PECAM-1 staining in E12.5 *Zmiz1*^fl/fl^ and *Zmiz1*^cECKO^ yolk sacs. Note the absence of distinct, organized vessels within the yolk sac of the mutants. Scale bars: 100 μm. Error bars represent mean ± s.e.m; two-tailed unpaired t-test. ns (not significant; P>0.05), *P<0.05, **P<0.01, ***P<0.001, ****P < 0.0001.

### Endothelial-Zmiz1 is required for postnatal retinal angiogenesis but not vascular maintenance

To elucidate the role of Zmiz1 in the endothelium during physiological postnatal angiogenesis, we generated inducible EC-specific Zmiz1 knockout mice. This was accomplished by crossing the *Zmiz1*^fl/fl^ line with mice carrying tamoxifen-inducible Cre recombinase under control of the EC-specific *Cdh5* promoter (Cdh5(PAC)-iCre^ERT2^)^28^. Cre-mediated inactivation of *Zmiz1* in ECs (*Zmiz1*^fl/fl^;*Cdh5*-Cre^ERT2^, referred to as *Zmiz1*^iECKO^; *Zmiz1*-inducible EC knockout mice) was induced by daily oral administration of tamoxifen from P1-P3 for early induction (Fig. 2A). Tamoxifen treatment resulted in an approximately 80% reduction of *Zmiz1* mRNA levels in isolated lung ECs (iLECs) of *Zmiz1*^iECKO^ pups compared to littermate controls (*Zmiz1*^fl/fl^) at P7 (Fig. 2B). Retinal blood vessels were analyzed at P7 or P10 following early induction, or at P12 following late induction. Whole-mount retinas immunofluorescently stained for the retinal endothelial cell marker, isolectinB4 (IB4), showed a significant reduction of vascular outgrowth, density, and branching in *Zmiz1*^iECKO^ as compared to littermate controls at P7 (Fig. 2C-F). Moreover, impaired retinal angiogenesis was also observed in the heterozygote mice (*Zmiz1*^fl/wt-iECKO^), which exhibited decreased vascular outgrowth and density in comparison to the P7 controls (Supplementary Fig. 2). However, the vascular defects were not as severe as those observed in the homozygous *Zmiz1*^iECKO^ mutants. Interestingly, *Zmiz1*^iECKO^ mice also exhibited defects in the overall number of retinal arteries and veins at P7. Immunofluorescent antibody staining of retinas with IB4 and alpha-SMOOTH MUSCLE ACTIN (α-SMA) revealed a decreased number of main arteries and veins in Zmiz1 mutant retinas as compared to *Zmiz1*^fl/fl^ mice (Fig. 2G-I; Supplementary Fig. 2). To further assess vascular morphology at later stages, P10 retinas from early deletion of *Zmiz1* (P1-P3) were analyzed (Fig. 2J). At this time point, the vasculature in the retina has typically reached the peripheral edge and superficial capillaries have started sprouting vertically to give rise to the deep vascular plexus in the outer plexiform layer. However, at P10, vascular outgrowth and perpendicular sprouting into the deeper plexus were strongly impaired in *Zmiz1*^iECKO^ mice versus littermate controls (Fig. 2K-L and data not shown). Thus, collectively, the early Zmiz1 deletion data demonstrated that endothelium-specific loss of Zmiz1 in postnatal mice leads to significant deficits in angiogenesis and an overall reduced vascular presence.

**Figure 2.**
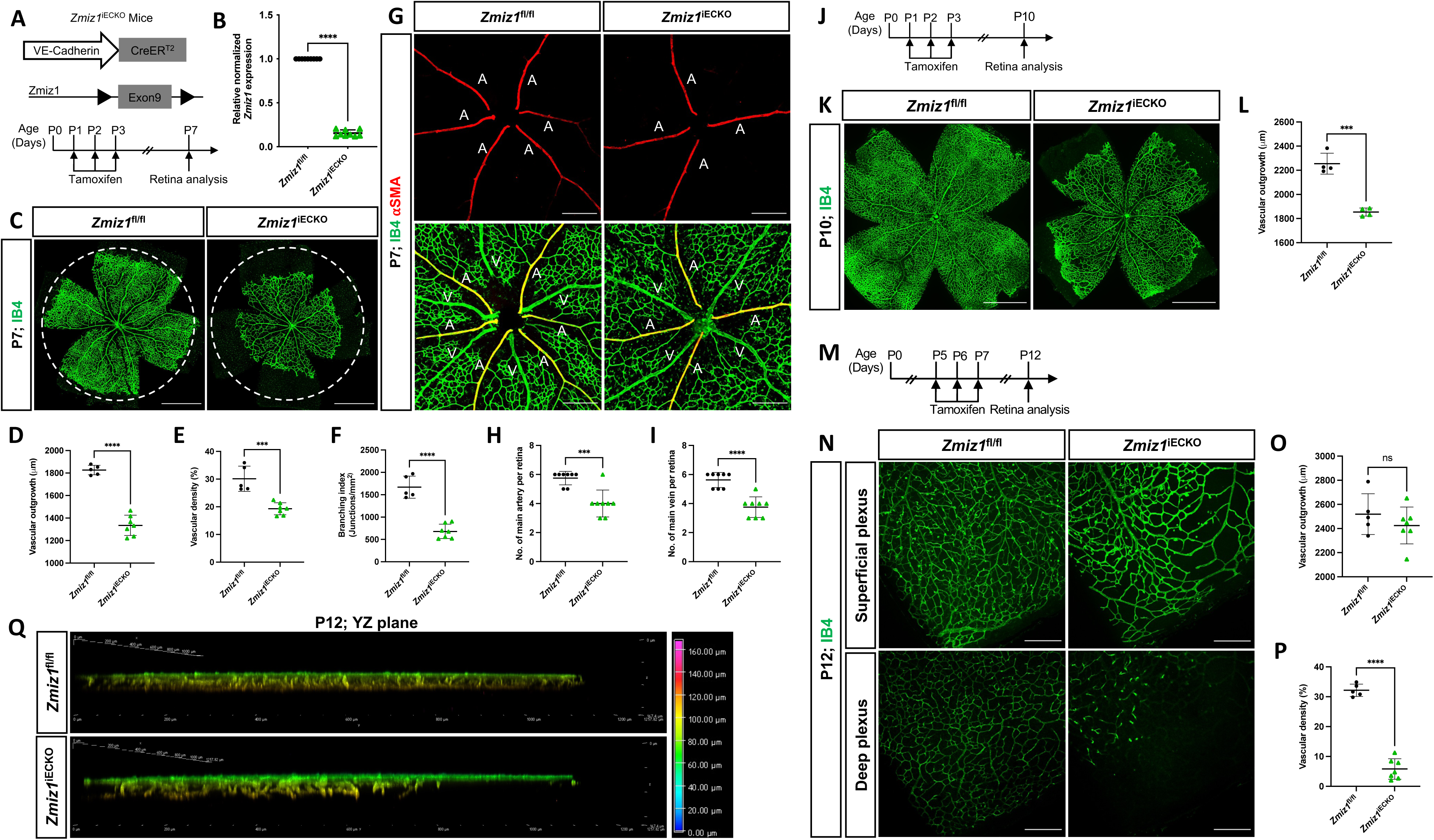
Zmiz1 EC deficiency leads to impaired retinal vascular morphogenesis. **A,** Schematic illustration of the VE-Cadherin (*Cdh5*)-Cre^ERT2^ transgene and Cre-mediated recombination of *Zmiz1* floxed (fl) exon 9 to generate EC-specific deletion of *Zmiz1* (*Zmiz1* induced endothelial cell knockout; *Zmiz1*^iECKO^). Time points utilized within the study for tamoxifen administration and retinal analysis are noted. **B,** Relative *Zmiz1* expression levels in isolated lung ECs from *Zmiz1*^fl/fl^ and *Zmiz1*^iECKO^ mice at P7 as determined via qRT-PCR (n=3). **C,** Whole-mount retinas stained for isolectin-B4 (IB4). (Dotted white circle represent outgrowth in *Zmiz1*^fl/fl^ retina). Scale bar, 1000 μm. **D-F,** Quantification of indicated morphometric parameters analyzed within *Zmiz1*^fl/fl^ (n=5) and *Zmiz1*^iECKO^ (n=7) retinas at P7. **G**, Whole-mount retinas stained for IB4 and alpha-SMOOTH MUSCLE ACTIN (α-SMA). Scale bar, 250 μm. **H** and **I,** Quantification of main arteries and veins within *Zmiz1*^fl/fl^ (n=8) and *Zmiz1*^iECKO^ (n=8) retinas at P7. **J,** Schematic showing tamoxifen administration in pups for early induction studies and analyzed at P10. **K,** Representative images of whole-mount retinas stained with IB4 at P10. Scale bars 1000 μm. **L,** Quantification of vascular outgrowth in *Zmiz1*^fl/fl^ (n=4) and *Zmiz1*^iECKO^ (n=4) retinas at P10. **M,** Schematic showing tamoxifen administration in pups for late induction studies. **N,** Representative images of IB4 vessels in superficial and deep layers at P12. Scale bars: 250 μm. **O** and **P,** Quantification of vascular outgrowth and deep vascular plexus density in *Zmiz1*^fl/fl^ (n=5) and *Zmiz1*^iECKO^ (n=7) retinas at P12. **Q,** 3D vertical view in the YZ plane with depth coding of *Zmiz1*^fl/fl^ and *Zmiz1*^iECKO^ retinas at P12. Note the absence of vertical sprouting in the periphery (to the right) of the *Zmiz1*^iECKO^ retinas. Error bars represent mean ± s.e.m; two-tailed unpaired t-test. ns (not significant; P>0.05), *P<0.05, **P<0.01, ***P<0.001, ****P < 0.0001.

These initial analyses primarily focused on the retina vasculature in a two-dimensional fashion (superficial vessels). Consequently, we evaluated the angiogenic role of Zmiz1 in three-dimensional space by analyzing deep vascular plexus formation in the retina. Introduction of tamoxifen from P5-P7, when many of the vessels have differentiated and patterned properly but the vasculature is still remodeling and migrating to the periphery (Fig. 2M), resulted in *Zmiz1*^iECKO^ retinas with a relatively normal patterned superficial vascular plexus that reached the periphery at P12 (Fig. 2N, O). However, the vascularization of the deeper vascular plexus was significantly affected, as notably fewer vessels were observed in comparison to *Zmiz1*^fl/fl^ mice (Fig. 2N-Q). Specifically, compared to control retinas, deep layer vessels were absent at the periphery in Zmiz1 mutants, while the vessels that did invade more centrally appeared disorganized (Fig. 2Q). Therefore, these findings indicated that Zmiz1 is also important for the perpendicular, angiogenic sprouting events required to populate the deeper layer of the retina.

Based on the findings above, Zmiz1 is essential during periods of active retinal angiogenesis, which is consistent with its expression levels in ECs during early mouse retina development^26^ (Supplementary Fig. 1D). However, the role of Zmiz1 in vascular maintenance during adulthood was unknown. To address this, we induced deletion of *Zmiz1* in adult mice from P28-P31 and analyzed retinas at P42 (Supplementary Fig. 3A). Interestingly, *Zmiz1*^iECKO^ mice did not display any obvious vascular patterning or morphological defects (Supplementary Fig. 3B). Evaluation of vascular density in both the superficial and deep vascular plexi revealed no significant difference in control and *Zmiz1*^iECKO^ mice (Supplementary Fig. 3C-E). Thus, these results suggested that Zmiz1 is not required for vascular maintenance during adulthood, which notedly corresponds to a continuous decrease in *Zmiz1* endothelial cell expression from P15-P50 in the retina^26^ (Supplementary Fig. 1D).

### Loss of Zmiz1 leads to increased retinal vessel regression

During postnatal retinal angiogenesis, the growth of initial blood vessels to the formation of a mature vascular network involves various cellular processes, such as EC migration, proliferation, apoptosis, vessel regression, and vessel remodeling. To assess whether some of these processes were defective and contributed to the vascular phenotypes in *Zmiz1*^iECKO^ retinas, we first analyzed EC proliferation and apoptosis. EC proliferation was assessed by co-staining for ETS-related gene (ERG; EC-specific nuclei marker) and the proliferation marker Ki67. We observed no significant difference in the rate of EC proliferation between *Zmiz1*^iECKO^ and control retinas at P7 (Fig. 3A, B). Similarly, co-staining for CLEAVED CASPASE 3 (a key protease in apoptosis) and ERG revealed that EC apoptosis at P7, which is normally low, was unaffected by loss of Zmiz1 (Fig. 3C, D). Next, we assessed blood vessel regression by co-immunolabeling for IB4 and the basement matrix component COLLAGEN IV. During vascular remodeling, COLLAGEN IV sleeves devoid of ECs (COLLAGEN IV positive; IB4 negative) are observed as vessels regress, and alterations in this process are typically associated with unstable, defective vascular remodeling. We identified a significant increase in the number of COLLAGEN IV sleeves within *Zmiz1*^iECKO^ retinas compared to the controls, indicative of increased vessel regression and decreased vascular stability (Fig. 3E, F). In addition, we examined pericyte vessel coverage since EC-pericyte interactions are also known to affect the angiogenesis process^29,30^. However, immunofluorescent antibody staining for pericyte marker NEURON-GLIAL ANTIGEN 2 (NG2) and IB4 revealed no noticeable differences in pericyte coverage between the control and *Zmiz1*^iECKO^ retinas at P7 (Fig. 3G, H). These data established that EC proliferation, EC apoptosis, and pericyte coverage are unchanged in the absence of Zmiz1. Conversely, loss of *Zmiz1* in ECs resulted in substantial increases in vessel regression indicating an overall contribution of Zmiz1 to vascular stability during angiogenic remodeling of the retinal vascular network.

**Figure 3.**
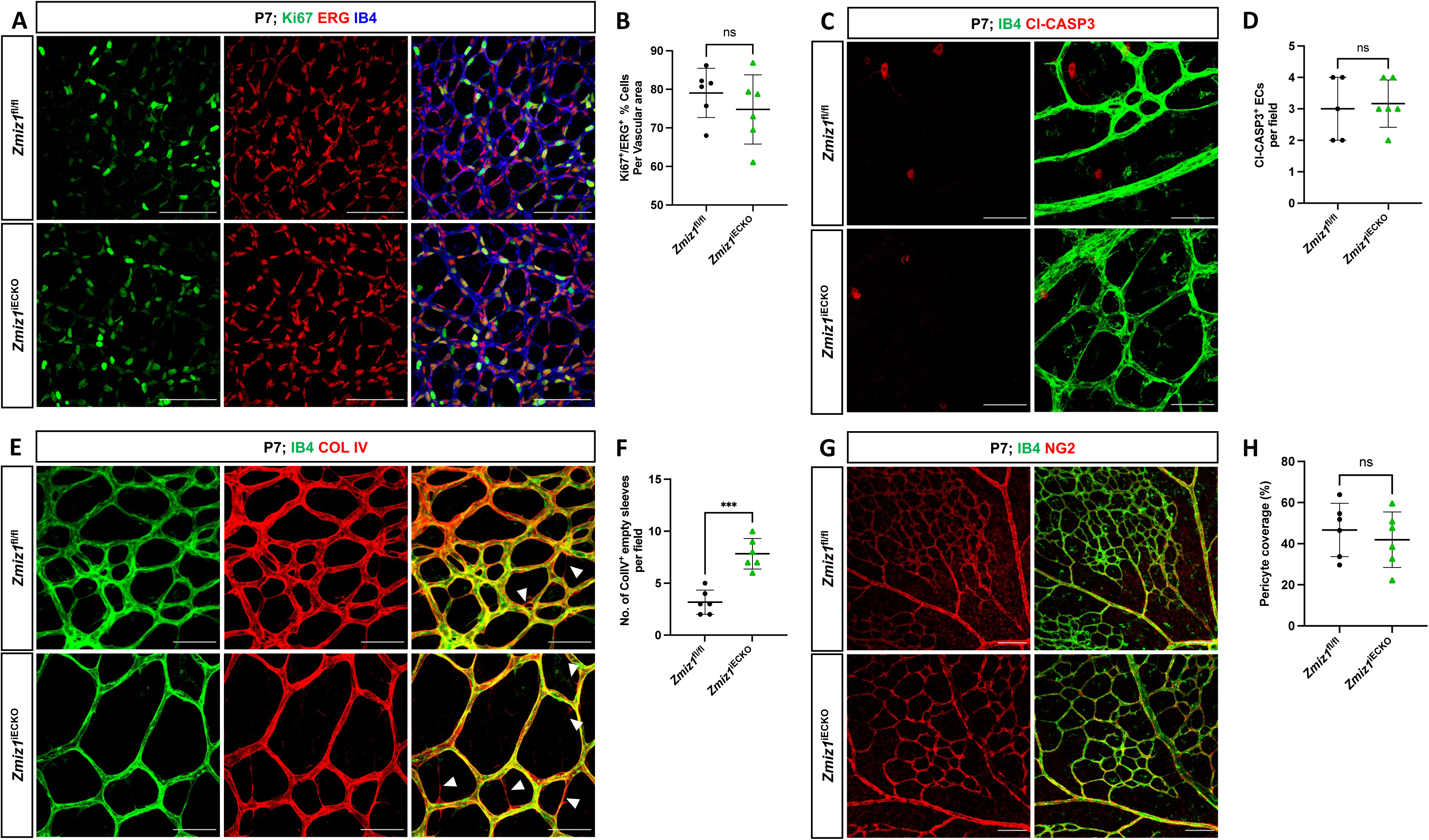
Deletion of Zmiz1 in ECs results in increased retinal vascular regression. **A,** Representative images of P7 retinas stained with Ki67, ERG, and IB4. Scale bars: 100 μm. **B,** Quantification of Ki67^+^ proliferating ECs in *Zmiz1*^fl/fl^ (n=6) and *Zmiz1*^iECKO^ (n=6) mice at P7. **C,** Representative images of P7 retinas stained with IB4 and CLEAVED CASPASE 3 (Cl-CASP3). Scale bars: 50 μm. **D,** Quantification of Cl-CASP3^+^ apoptotic ECs in *Zmiz1*^fl/fl^ (n=6) and *Zmiz1*^iECKO^ (n=6) mice at P7. **E,** Representative images of P7 retinas stained with IB4 and COLLAGEN IV (white arrowheads indicate empty COLLAGEN IV sleeves devoid of ECs). Scale bars 50 μm. **F,** Quantification of empty Collagen IV sleeves in *Zmiz1*^fl/fl^ (n=6) and *Zmiz1*^iECKO^ (n=6) retinas at P7. **G,** Representative images of P7 retinas stained with IB4 and NEURAL/GLIAL ANTIGEN 2 (NG2). Scale bars 100 μm. **H,** Quantification of pericyte coverage in *Zmiz1*^fl/fl^ (n=6) and *Zmiz1*^iECKO^ (n=6) retinas at P7. Error bars represent mean ± s.e.m; two-tailed unpaired t-test. ns (not significant; P>0.05), ***P<0.001.

### Zmiz1 mutants exhibit defective sprouting angiogenesis

During initial observations of the retinal vasculature (Fig 2), we noticed an obvious difference in the sprouting angiogenic blood vessels at the vascular front between Zmiz1 control and mutant retinas. Specifically, there appeared to be fewer vessels sprouting toward the retina periphery in *Zmiz1*^iECKO^ retinas. Indeed, quantification confirmed a significant decrease in the number of P7 vascular sprouts in Zmiz1 mutant retinas compared to controls (Fig. 4A, B). This reduction was notable because deficits in sprouting angiogenesis regularly result in retinal vascular phenotypes like those observed in the *Zmiz1*^iECKO^ retinas^31,32^. Therefore, to further evaluate the role of Zmiz1 in sprouting angiogenesis, we performed the fibrin bead assay with immortalized human aortic ECs (telomerase human aortic EC; TeloHAEC). Beads were coated with equal numbers of TeloHAECs treated with either scramble-control siRNA or *Zmiz1*-targeted siRNA and allowed to sprout for 120 hours. Control siRNA treated TeloHAECs showed robust sprout formation, whereas cells subjected to *Zmiz1* siRNA displayed a significant reduction in the length and number of sprouting vessels (Fig 4C, D). Quantitative polymerase chain reaction (qPCR) verified that *Zmiz1* expression was appreciably reduced in *Zmiz1* siRNA TeloHAECs (Fig. 4E). Additionally, fibrin bead assays were carried out using a mouse immortalized EC line (MS1) and resulted in a similar decrease in sprout formation when Zmiz1 was depleted (Supplementary Fig. 4A, B); reduction in the number of sprouts was not due to changes in EC proliferation *in vitro* (Supplementary Fig. 4C, D). Since vascular endothelial growth factor-A (VEGF-A) is a major pro-angiogenic factor involved in sprouting angiogenesis^33–35^, we also tested whether Zmiz1 might be involved in VEGF-A-mediated sprout formation. We conducted TeloHAEC bead assays in the presence of VEGF-A and compared them to our previous studies. As expected, we found that sprouting vessel numbers substantially increased in control siRNA treated ECs when VEGF-A was present versus absent (Fig. 4C, D; gray bars). Similarly, *Zmiz1* siRNA-treated TeloHAECs exhibited more sprouts in the presence of VEGF-A as compared to those lacking VEGF-A (Fig. 4C, D; red bars). However, the differences in the number of sprouts between the *Zmiz1* siRNA group with (red bar) and without (gray bar) VEGF-A (∼2.35 sprouts; *P<0.05) compared to the control siRNA group plus (red bar) or minus (grey bar) VEGF-A (6.9 sprouts; ****P < 0.0001) were not as significant indicating Zmiz1 participation in VEGF-A mediated sprout formation. Moreover, the reduction in the number of sprouting vessels when *Zmiz1* siRNA treatments were compared to corresponding controls (control versus *Zmiz1* siRNA TeloHAEC [gray bars] and control versus *Zmiz1* siRNA TeloHAEC with VEGF-A [red bars]) showed significant differences as well. Quantification indicated a reduction of approximately 7.6 sprouts in the absence of VEGF-A, but a reduction of approximately 12 sprouts when VEGF-A was present (Fig. 4D), further supporting the notion that VEGF-A-mediated sprout formation requires Zmiz1.

**Figure 4.**
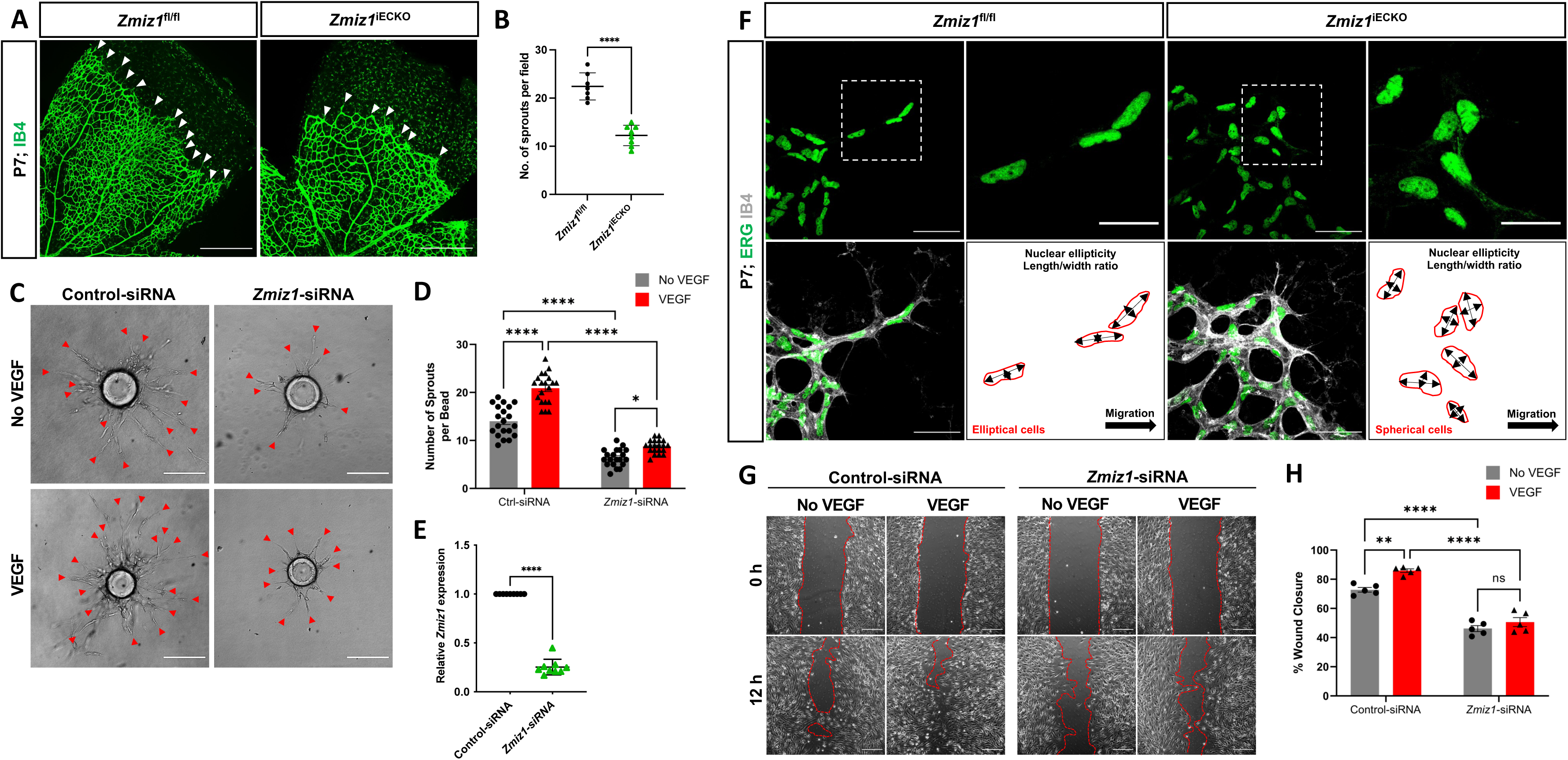
Zmiz1 is required for sprouting angiogenesis and EC migration. **A,** Images of IB4 immunolabeled retinas at the vascular front in P7 *Zmiz1*^fl/fl^ and *Zmiz1*^iECKO^ mice (white arrowheads mark the leading-edge vascular sprouts). Scale bar, 250 μm. **B,** Quantification of the number of sprouts within *Zmiz1*^fl/fl^ (n=7) and *Zmiz1*^iECKO^ (n=8) retinas at P7 (quantifications represent the average number of sprouts within each leaflet). **C,** Representative images of bead assays embedded in 3D fibrinogen gel at 120 hours following control and *Zmiz1* siRNA treatments and in the presence or absence of VEGF-A (red arrowheads mark sprouts emanating from the bead). Scale bar, 100μm**. D,** Quantification of the number of sprouts per bead (n=20). **E,** qRT-PCR analysis of control and Zmiz1 siRNA treated TeloHAECs for *Zmiz1* mRNA levels normalized to *GAPDH* transcripts (n=3). **F,** Close-up images of retinas stained for IB4 and ETS related gene (ERG) to mark EC nuclei at the vascular front within P7 *Zmiz1*^fl/fl^ (n=7) and *Zmiz1*^iECKO^ (n=8) retinas. Cell-shape schematics indicating nuclear ellipticity within each group are represented. Scale bar, 50 μm and 25 μm (insets highlight sprouts). **G,** Images of scratch wound assays performed on TeloHAEC confluent monolayers following control and *Zmiz1* siRNA treatments and in the presence or absence of VEGF-A. Images at 0 and 12 hours following the scratch are depicted. Scale bar, 200μm. **H,** Quantification of the percentage of wound closure in TeloHAECs subjected to the various treatments in **G** (n=5). Error bars represent mean ± s.e.m; two-way ANOVA: *P<0.05, **P<0.01, ***P<0.001, ****P < 0.0001.

Previous reports noted defective migration and polarization of neuronal cells upon loss of Zmiz1 function *in vivo*^16^. This led us to explore the possible role of Zmiz1 in the regulation of tip-cell behavior and migration during sprout formation/elongation. In this context, we first analyzed the nuclei shape of tip-cell-associated ECs at the vascular front; elliptical nuclei are often connected with directed migration, while spherical-shaped nuclei are linked to non-migratory cells^36^. In control mice, ERG-positive nuclei of tip ECs were predominately elliptical in shape and directed toward the avascular area where the retinal vessels typically migrate. (Fig. 4F). However, in *Zmiz1*^iECKO^ mice, the nuclei of ECs at the leading edge were more spherical, including those in the comparatively shorter sprouts, suggesting an impaired cell migration phenotype (Fig. 4F). To obtain a better grasp on Zmiz1 function in EC migration, cell migration assays (scratch assay) were carried out with TeloHAECs. Under normal culture conditions, *Zmiz1* siRNA TeloHAECs showed a decreased repopulation of cells into the “wound” area in comparison to control siRNA-treated ECs suggesting defective migratory properties in the absence of Zmiz1 (Fig. 4G, H). Upon VEGF-A stimulation, control siRNA TeloHAECs displayed enhanced migration and closure of the scratch area compared to ECs in the normal culture conditions as expected. However, the *Zmiz1* siRNA-treated group showed no significant wound closure whether VEGF-A was present or absent, further underscoring that Zmiz1 is necessary to propagate the pro-migratory effects of VEGF-A in this *in vitro* assay.

### Zmiz1 inactivation leads to gene expression changes in the endothelium, including reduced expression of tip-cell-associated genes

To further define the role of Zmiz1 in the regulation of EC behavior, we performed transcriptional profiling of isolated retinal ECs (iRECs) at P7 (Fig. 5A). RNA-Sequencing (RNA-seq) analysis of three biological replicates of iRECs from control and *Zmiz1*^iECKO^ mice revealed 854 upregulated and 1530 downregulated genes following the loss of *Zmiz1* (Fig. 5B). Gene ontology (GO) analysis unveiled upregulated genes enriched for biological processes such as immune response and T-cell activation (Fig. 5C), which is relevant to previous work demonstrating an important role for Zmiz1 in T cell development and leukemogenesis^11^. Importantly, downregulated genes, representing targets that are likely positively regulated by Zmiz1, were enriched for GO terms associated with developmental processes such as angiogenesis, blood vessel morphogenesis and vascular development (Fig. 5D). These transcriptional findings were in alignment with our phenotypic embryonic and retinal studies (Fig 1-4). As an additional measure, we performed RNA-seq analysis on control-shRNA and *Zmiz1*-shRNA treated MS1 ECs (Supplementary Fig. 5). Deletion of Zmiz1 resulted in 1711 differentially expressed genes, of which 718 genes were upregulated while 993 genes were downregulated (Supplementary Fig. 5A). Notably, GO analysis of the top downregulated genes were associated with EC migration, angiogenesis, vascular development, and blood vessel morphology, similar to the iREC RNA-seq results (Supplementary Fig. 5B, C). Intriguingly, further assessment of the MS1 RNA-seq data implicated differentially expressed genes that were targets of several Notch-associated transcription factors, including Hes Related Family BHLH Transcription Factor With YRPW Motif 1 (Hey 1), Hairy/Enhancer of Split Related Protein 1 (Hes1), Notch 2 and 3 (Supplementary Fig. 5B, C); previous works showed that Zmiz1 transcriptionally works in coordination with Notch1 and Recombination Signal Binding Protein For Immunoglobulin Kappa J (RBPJ) in cancer and T-cell development^11,37^

**Figure 5.**
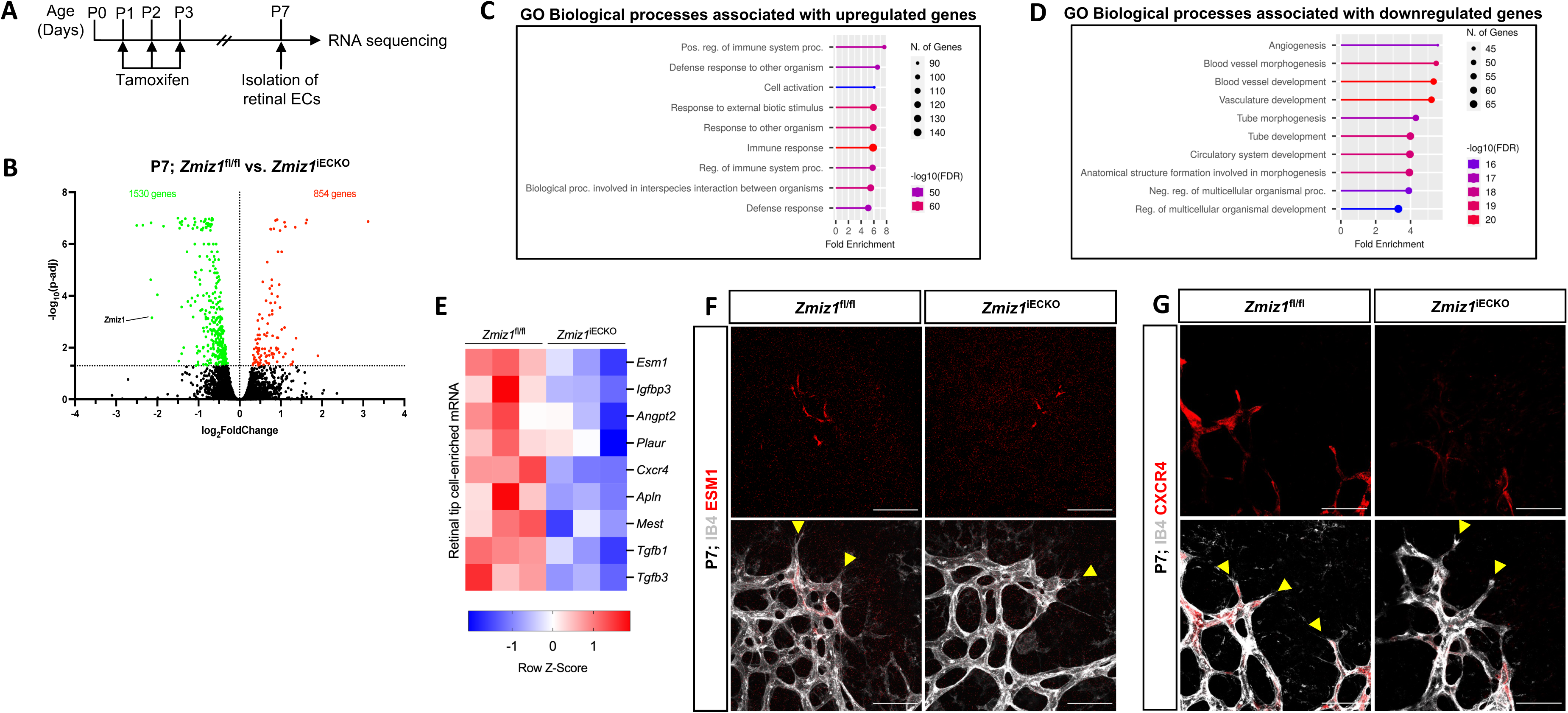
Expression of retinal endothelial tip-cell markers is downregulated upon Zmiz1 genetic ablation. **A,** Illustration depicting the strategy used for RNA-seq analysis on isolated retinal ECs (iRECs). **B,** MA plot of differentially expressed genes between *Zmiz1*^fl/fl^ and *Zmiz1*^iECKO^ iRECs at P7. Statistically significant upregulated and downregulated genes are depicted as red and green dots, respectively. Zmiz1 is highlighted and confirms reduced expression in *Zmiz1*^iECKO^ iRECs. **C** and **D,** Gene ontology (GO) analysis of biological process terms enriched in upregulated (**C**) and downregulated (**D**) genes of *Zmiz1*^iECKO^ iRECs (false discovery rates (FDR) are shown). **E,** Clustered heat map of gene count Z scores for retinal tip-cell enriched mRNAs. Columns represent pools of different biological samples. **F** and **G,** Close-up views of *Zmiz1*^fl/fl^ and *Zmiz1*^iECKO^ retinal tip-cell regions immunofluorescently labeled for IB4 and the tip-cell enriched markers ESM1 (**F**) and CXCR4 (**G**). Scale bars, 25μm. Notice overall reduced and sometimes absent expression of ESM1 and CXCR4 in the tip-cells (yellow arrowheads) of Zmiz1 retinas.

Given our overall findings connecting Zmiz1 function to pro-angiogenic growth, especially at the retinal vascular front, we examined the iREC RNA-seq data for indications that Zmiz1 affects transcriptional regulation of tip-cell-associated genes involved in retinal angiogenesis. Analysis revealed that several notably well-characterized genes either enriched in tip-cell expression and/or involved in tip-cell formation and function, such as Apln, Angpt2, Esm1, and Cxcr4 were markedly downregulated in Zmiz1 mutant retinas (Fig. 5E). Furthermore, immunofluorescent antibody stains for ESM1 and CXCR4 confirmed that expression of these tip-cell markers was substantially reduced at the vascular front in Zmiz1 deficient retinas, while controls showed typical, strong expression in ECs at the leading edge (Fig. 5F, G). Together, the iREC RNA-seq data and expression analysis provide further evidence that Zmiz1 is a critical tip-cell regulator required for proper sprouting angiogenesis.

### Loss of Zmiz1 in an OIR retinopathy model leads to reduced neovascularization

Zmiz1 has been linked to various pathological features and diseases, such as leukemia^11^, osteosarcoma^13^, diabetes^14^, erythropoiesis^12^, and several neurodevelopmental disorders^16–18^. To investigate whether Zmiz1 might have a role in pathological angiogenesis, we performed oxygen-induced retinopathy (OIR) studies, which are designed to emulate conditions similar to ocular retinopathies. Both Zmiz1 control and mutant pups were housed in a hyperoxia chamber (75% oxygen) from P7 to P12 to allow blood vessel regression and death. At P12 they were transferred to normal, room oxygen levels to create a hypoxic environment that promotes neovascularization (Fig. 6A). Pups were induced with tamoxifen for 3 consecutive days once placed in a normal oxygen environment (P12-14) and retinas were harvested and analyzed at P17. Compared to the *Zmiz1*^fl/fl^ mice, Zmiz1 deficient retinas exhibited significantly larger areas of vaso-obliteration (areas devoid of vasculature) at P17 (Fig. 6B-D). Accordingly, *Zmiz1*^iECKO^ retinas displayed significantly fewer regions with neovascular tufts than *Zmiz1*^fl/fl^ retinas (Fig. 6B-D). Neovascular tufts are areas of active growth and revascularization that occur in the OIR retinal model as the blood vessel network undergoes a corrective remodeling phase. Hence, these experimental results demonstrated that Zmiz1 is important for hypoxia-regulated angiogenesis during the revascularization process in a pathological setting.

**Figure 6.**
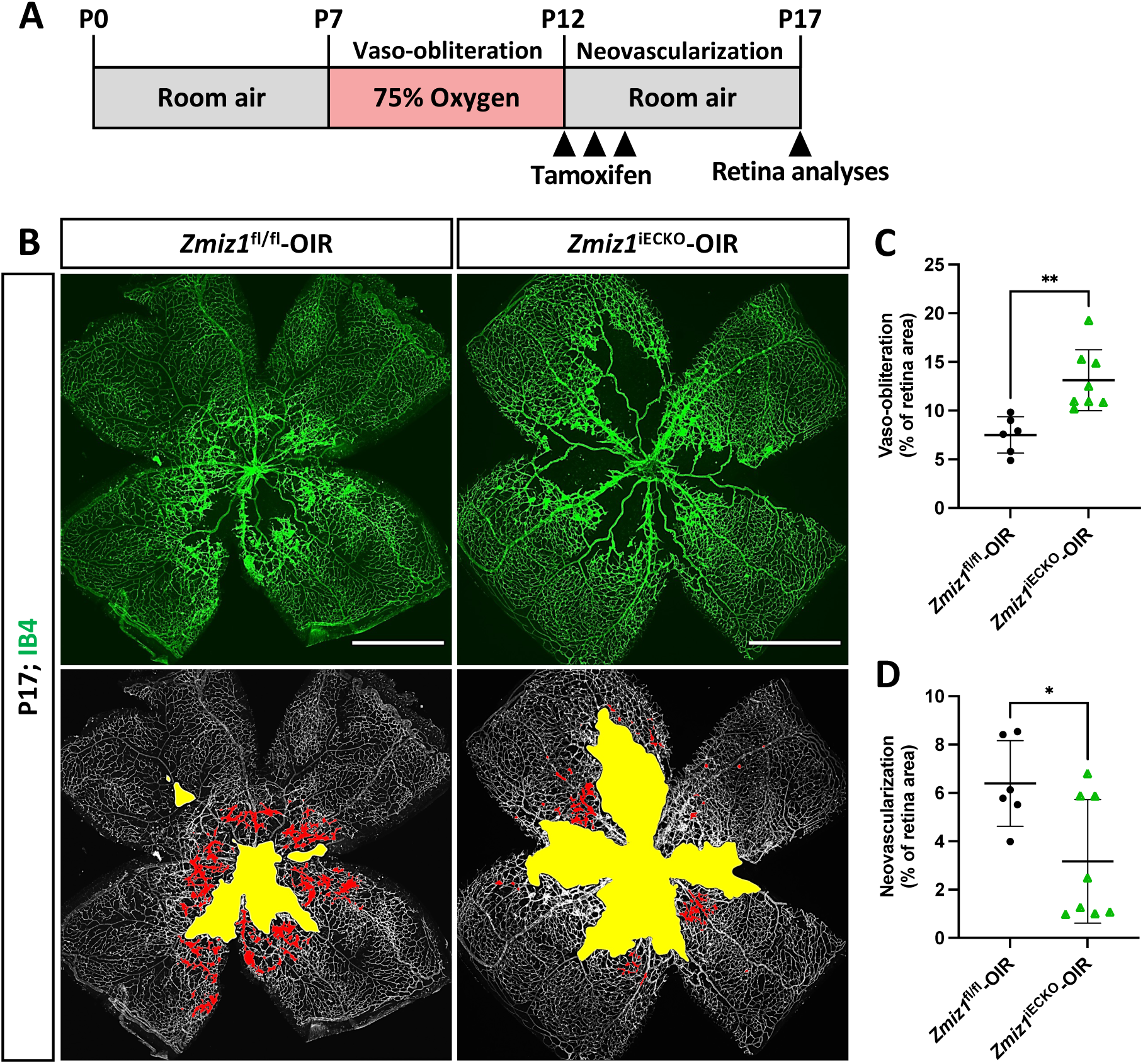
Loss of endothelial Zmiz1 impairs pathological angiogenesis. **A,** Schematic summary of the OIR strategy implemented to assess Zmiz1 in pathological neovascularization. **B,** Representative whole-mount images of the vasculature immunolabeled with IB4 in *Zmiz1*^fl/fl^-OIR (n=6) and *Zmiz1*^iECKO^-OIR (n=8) P17 retinas. Yellow space demarcates the avascular area, while the red markings highlight the neovascular tufts within each retina. Scale bars: 1000 μm. **C** and **D,** Quantification of the areas of vaso-obliteration (avascular space) and retinal neovascularization in *Zmiz1*^fl/fl^-OIR (n=6) and *Zmiz1*^iECKO^-OIR (n=8) P17 mice. Error bars represent mean ± s.e.m; two-tailed unpaired t-test. ns (not significant; P>0.05), *P<0.05, **P<0.01, ***P<0.001, ****P < 0.0001.

### Summary

The process of new blood vessel formation is vital for organismal growth and survival. Our study highlighted an important role of Zmiz1 during embryonic and postnatal vascular development. We showed that constitutive loss of endothelial *Zmiz1* during embryogenesis resulted in vascular defects and embryonic lethality. Using the inducible EC-specific Cre mice, we demonstrated that sprouting angiogenesis was disrupted in Zmiz1-deficient ECs both in physiological and pathological setting, providing evidence of Zmiz1 as an important regulator of the developing vasculature. Specifically, early postnatal deletion of endothelial *Zmiz1* resulted in delayed retinal outgrowth of superficial vascular plexus both at P7 and P10. In addition, late induction of endothelial *Zmiz1* deletion severely impaired perpendicular vascular sprouting resulting in reduced densities of vascular plexus in the deep layer at P12. The phenotype of disrupted vascular outgrowth in *Zmiz1* mutants is potentially attributed to the reduction in tip cell numbers and defective morphology, subsequent decreased sprout formation and reduced expression of crucial tip-cell genes. We also observed that *Zmiz1* is critical for intraretinal vascularization following ischemic injury in the OIR retina.

Although our work provides new information detailing the role of Zmiz1 in angiogenesis, a full appreciation of the mechanisms by which *Zmiz1* regulates blood vessel growth via angiogenesis will require additional investigation. However, transcriptional profiling of *Zmiz1*-deficient ECs in this study did uncover some interesting targets that are potentially regulated by Zmiz1. In particular, we found genes associated with acquisition of specialized tip/stalk-cell phenotypes in ECs during the sprouting process. The VEGF-A and NOTCH signaling pathways are critical players regulating tip-cell and stalk-cell selection whose ratio is critical during the angiogenic sprouting process in physiological and pathological conditions^38,39^. It is possible that the defective patterning of arteries observed in *Zmiz1*^iECKO^ mice is due to misregulation of DLL4-NOTCH1 pathway as this signaling cascade is important for arteriogenesis during vascular development^40^. This is further supported by previous work showing Zmiz1 has been linked with the NOTCH signaling pathway, specifically NOTCH1 in regulation of T-cell phenotype in Leukemia^11^. Moreover, our *in vitro* transcriptomic studies suggested that several differentially expressed genes misregulated in the absence of Zmiz1 are Notch TF targets (Supplementary Fig. 5). Lastly, microarray analysis following inactivation of VEGF-A in cultured human umbilical vein ECs (HUVECs) revealed downregulation of Zmiz1 in the blood-vessel gene cluster^20^. Taken together these studies indicate a potential link between VEGF-A, NOTCH1 and Zmiz1 in regulation of the angiogenesis process which remains to be fully elucidated. Furthermore, we acknowledge that Zmiz1 may regulate angiogenesis *in vivo* through additional mechanisms that are independent of the VEGFA and NOTCH signaling pathways.

Our study also identified that the loss of Zmiz1 in ECs influenced early vascular development, as evidenced by the aberrant embryonic vasculature and inhibited vascular expansion and impaired perpendicular branching of retinal vasculature. However, Zmiz1’s inactivation in mature vasculature with quiescent ECs had no effect on overall blood vessel integrity and remodeling. These findings suggest a clearly defined role of Zmiz1 in early embryonic and postnatal life, which will be essential to unraveling its molecular regulatory mechanisms. Interestingly, gene set enrichment analysis (GSEA) of gene sets identified during different developmental stages revealed higher enrichment of genes associated with NOTCH signaling pathway at P6^26^. Thus, considering the connection of Zmiz1 to NOTCH signaling and the similar developmental timing, it seems likely that Zmiz1 interacts with key putative TFs regulating EC behavior during early phase of vascular development.

In the field of vascular biology, many TFs that are important for EC differentiation and maturation have been identified, however their mechanisms of regulation are not fully defined. This study introduces a transcription cofactor with a previously unknown role in vascular development and function that may be involved in regulating the activity of vascular transcription factors, including those described above. Thus, our study delineates an important role of endothelial *Zmiz1* in physiological and pathological angiogenesis.

## Methods

### Mice and breeding

All the mice were housed in individual ventilated cages, in a temperature-controlled room with a 12 h light/dark cycle. All animal protocols in this study were performed in accordance with Tulane University’s Institutional Animal Care and Use Committee policies. In our studies the following transgenic mouse lines were used: *Zmiz1*^fl/fl^, *Tie2*-Cre, and *Cdh5*-Cre^ERT2 41,42^. All mice were maintained in a mixed genetic background. *Zmiz1*^fl/fl^ mice were crossed with both *Tie2*-Cre and *Cdh5*-CreERT2 lines for embryonic and postnatal studies respectively. Littermate controls of both sexes were used in all experiments. Genotyping was performed as previously detailed^43^.

### Tamoxifen treatment

To induce Cre activity, 100μg of tamoxifen (Sigma, T5648) was administered orally to newborn pups born from *Zmiz1*^fl/fl^;*Cdh5*-Cre^ERT2^ and *Zmiz1*^fl/fl^ mating pairs from P1-P3 for early deletion or from P5-P7 for late deletion. For 4-week-old mice, 2 mg of tamoxifen was injected intraperitoneally daily from P28-P31.

### Embryo dissection, processing, and immunofluorescent analysis

Embryos were collected from pregnant females at indicated time points following timed mating, designating embryonic day 0.5 (E0.5) as noon on the day a vaginal plug was observed. Embryos with yolk sacs were dissected and imaged by brightfield in 1X PBS, followed by overnight fixation in 4% PFA at 4°C. Next, both yolk sacs and embryos were incubated in PBST buffer (PBS with 0.5%Ttriton X-100) overnight at 4°C and then transferred into CAS-Block (Life technologies, 008120) for 4 h at room temperature. Both yolk sac tissues and whole embryos were incubated in primary antibody to rat anti-PECAM-1(BD Pharmingen, 553370) diluted in CAS-Block overnight at 4°C. Following primary antibody incubation, both yolk sacs and embryos were incubated overnight at 4°C with secondary antibodies to chicken anti-rat Alexa Fluor 488 (Life technologies, A21470) diluted in CAS-Block. Lastly, both yolk sacs and embryos were washed in PBST buffer before imaging. The amniotic membrane was used for genotyping purposes.

### Retina dissection, processing, and immunofluorescent analysis

Eyes were collected at P7, P10, P12, and P42 for analysis. Retinas were dissected out and whole-mount retina staining was performed as previously described^44^. Primary antibodies used for whole mount retina immunofluorescent staining: IB4-488 (1:250, Invitrogen, 121411), IB4-594 (1:250, Invitrogen, 12143), SMA-Cy3 (1:250, Sigma Aldrich, C6198), ERG-488 (1:250, Abcam, Ab196374), ERG-647 (1:250, Abcam, Ab196149), COLLAGEN IV (1:100, Millipore, AB756P), NG2 (1:100, Millipore, AB5320), Ki67-488 (1:250, Cell signaling, 11882), CL-CASPASE3 (1:100, Cell signaling, 9661), ESM-1 (1:100, R&D systems, AF1999), CXCR4 (1:100, R&D systems, MAB 21651), and ZMIZ1 (1:100, Santa Cruz, Sc-376825).

### Analysis of retinal vasculature

Quantifications of retinal vasculature were performed on high-resolution confocal images using Nikon NIS-elements and Angiotool analysis software^45^. Radial outgrowth was determined as the distance between the vascular front and the central optic nerve in each leaflet of the retina and averaged. Whole retinal images were used to measure the vascular density and the number of branching points. Quantification of sprouts was performed at the vascular front in 20X images.

### Isolation of murine retina and lung endothelial cells

Retinal ECs were isolated from P7 pups as previously described^46^. Briefly, isolation of ECs from the retinas was performed using the Miltenyi Neural tissue Dissociation Kit (P) with minor modifications (Miltenyi, 130-092-628). For each sample, 8-10 retinas were pooled and digested in enzyme mix 1 and enzyme mix 2 to obtain a single cell suspension. Cells were incubated with CD31 conjugated dynabeads for 30 min at 4°C. Following the incubation, CD31+ cells were separated from the cell mixture via magnetic activated cell sorting. Next, RNA was isolated from the cells and used for downstream RNA-sequencing. Lung ECs were isolated from P7 pups as previously described^44^. Lung ECs were isolated from both control and *Zmiz1*^iECKO^ mice and RNA was extracted for downstream assays.

### Cell culture and siRNA transfection

MS1 mouse pancreas ECs (CRL-2279; ATCC) and 293T cells (CRL-3216; ATCC) were cultured in DMEM medium supplemented with 5% fetal bovine serum (FBS) and Antibiotic-Antimycotic (Thermofisher, 15240062). TeloHAECs (ATCC, CRL-4052) were cultured in EBM-2 media (Lonza, CC-3156) supplemented with EGM-2 bullet kit (Lonza, CC-3162) according to manufacturer’s instructions. Human lung fibroblasts (NHLF) (Lonza, CC-2512) were cultured in fibroblast growth media supplemented with FGM-2 bullet kit (Lonza, CC-3132) All cells were maintained at 37°C and 5% CO_2_. TeloHAECs were seeded in a 6-well plate overnight. The following day, cells were transfected with 20μM of either control-siRNA (Dharmacon, D-001810-01-05) or *Zmiz1*-siRNA (Dharmacon, L-007034-00-0005) using Lipofectamine3000 (Thermofisher, L3000015) following manufacturer’s instructions.

### Lentivirus production and transduction

To generate lentiviruses, 293T cells were transfected with 2 μg of pMD2.G (Addgene, 12259), 4 μg of psPAX2 (Addgene, 12260), and 6 μg of GIPZ Non-silencing Lentiviral shRNA (control) and GIPZ Zmiz1 lentiviral shRNA (Dharmacon, RMM4532-EG328365) using 40 μl Lipofectamine LTX (Thermofisher, 15338100) in 500 µl Opti-MEM (Thermofisher, 31985062) per dish. After 24 h virus-containing supernatants were collected, centrifuged, filtered through a 0.45µm syringe filter (Thermofisher, F2500-1) and then stored at −80 °C until use. MS1 ECs were then transduced with the lentivirus and selected in puromycin (Thermofisher, A1113803) to generate (control-shRNA or *Zmiz1*-shRNA) stable cell lines.

### RNA extraction and quantitative RT-PCR

RNA was extracted from MS1 ECs, teloHAECs and iLECs using the GeneJET RNA Purification Kit (Thermofisher, K0732). First strand cDNA synthesis was performed using the iScript Reverse Transcription Supermix kit (Biorad, 1708840). Quantitative RT-PCR was carried out using the PerfeCTa SYBR Green Fastmix (Quantabio, 95071) and gene-specific primers (Table 1) on CFX96 system (Biorad). Relative gene expression was determined using the ΔΔCt method.

### Scratch assay

Scratch assay was performed on confluent control-shRNA and *Zmiz1*-shRNA stable MS1 ECs. A horizontal and vertical wound was generated in each well using a 200 μl pipet tip followed by three washes with PBS. Phase contrast images were taken right after the scratch (0 h) and 16 h after incubation at 37°C and 5% CO_2_. The percentage of wound closure was determined using the ImageJ software. For VEGF stimulation studies, confluent control-siRNA and *Zmiz1*-siRNA treated teloHAECs were serum starved overnight. A wound was introduced as described above and cells were stimulated with or without VEGF (50 ng/mL). Cell imaging and analysis were performed as described above.

### Fibrin gel bead assay

Fibrin bead sprouting assay was performed as previously described^46^. Briefly, control and *Zmiz1* siRNA treated teloHAECs, as well as control-shRNA and *Zmiz1*-shRNA MS1 ECs, were coated on cytodex beads and embedded into a fibrin gel. Fibroblasts were cultured on the gels and media was replaced every other day. Sprouting activity was monitored over the next few days and on day 5 the beads were imaged. Approximately 15 beads per condition were quantified for the number of sprouts per bead from 4 independent experiments. For VEGF stimulation studies, fibrin gel bead assays were similarly set up as described above. Cells were serum starved overnight and the next day stimulated with or without VEGF (50 ng/mL). Sprout imaging and analysis were performed as described above.

### *In vitro* cell proliferation assay

*In vitro* cell proliferation analysis was performed via the Click-iT EdU (5-ethynyl-2’-deoxyuridine) cell proliferation kit following manufacturer’s instructions. Briefly, Non-silencing and *Zmiz1*-shRNA MS1 ECs were incubated in EdU solution for 4h at 37°C and 5% CO_2_. Cells were then fixed and permeabilized, and EdU was detected via immunofluorescence analysis.

### RNA sequencing and differential gene expression analysis

RNA sequencing and gene expression analysis was performed as previously described^46^. Briefly, Total RNA extracted from iRECs and MS1 ECs was quantified and verified before library preparation for sequencing. Sequenced reads were aligned to the mouse (mm10) reference genome in the Basespace sequence hub (Illumina), and the aligned reads were used to quantify mRNA expression to determine differentially expressed genes. Gene ontology (GO) analysis of differentially expressed genes was performed using graphical gene-set enrichment tool for plants and animals—Shiny Go version 0.76^47^. Sequencing data were deposited in the Gene Expression Omnibus (GEO) database with accession code.

### OIR model

OIR studies were performed similar to previous work^46^. Neonatal pups and their nursing mother were exposed to 75% oxygen from P7-P12 in a designed chamber (Biospherix, ProOx110); from P12-P17 they were moved to room air^48^. Tamoxifen was administered to the pups orally from P12-P14. Vascular loss and neovascularization were quantified in P17 retinas using an automated OIR retinal image analysis software^49^.

### Statistical analysis

Data analysis was performed using Graphpad Prism 9.3.1. All values are presented as bar graphs with mean ± standard deviation shown as error bars. Statistical significance was determined by the two-tailed unpaired t-test between two conditions with a p-value less than or equal to 0.05 was considered statistically significant.

## Acknowledgements

We would like to thank Jovanny Zabaleta and Jone Garai at The Louisiana Cancer Research Consortium (LCRC) Translational Genomics Core Center for their continued support of our RNA-Seq experiments. This work was supported by the Tulane University Committee of Research (COR) Fellowship, NIH-R01 HL139713 (SMM), NIH-R01 HL163196 (SMM) and NIH/NIAID-R01 AI136941(MYC).

## Author Contributions

NRP and SMM developed and designed the experiments. NRP and RKC performed the experiments. NRP and SMM wrote the manuscript. RKC provided edits to the manuscript. MYC provided critical transgenic animals and edits to the manuscript.

## Notes

### Competing Interest Statement

The authors have declared no competing interest.

