## Supplemental Material for "Endothelial Zmiz1 modulates physiological and pathophysiological angiogenesis during retinal development"

**Supplementary figure legend:**

**Figure 1. Summary of EC-specific Zmiz1 expression within various tissues. A,** Analysis of *Zmiz1* expression levels in ECs and total tissue of P7 brain, kidney, liver, lung, and adult brain using the Vascular Endothelial Cell Trans-omics Resource Database (VECTRDB). **B,** *Zmiz1* expression in each cell type isolated from an adult mouse brain can be visualized within the violin plots (searchable database Tabula Muris https://tabula-muris.ds.czbiohub.org/)[16]. **C,** Analysis of *Zmiz1* mRNA expression levels in P7 isolated brain ECs and its vessel subtypes using single cell RNA-seq data in VECTRDB. **D,** Expression levels of *Zmiz1* in murine retinal ECs during postnatal development determined by available bulk RNA sequencing data (ref). **E,** Representative images of P7 retinas immunofluorescently labeled with IB4 and ZMIZ1 antibodies (A, artery; V, vein). Scale bars: 200 μm.

**Figure 2. Heterozygous loss of *Zmiz1* in ECs leads to defects in radial outgrowth and vascular density. A,** Representative fluorescent images of whole-mount retinas immunolabeled with antibodies against IB4 and α-SMA at P7. Scale bars: 1000 μm. **B** and **C,** Quantification of vascular outgrowth and vascular density in $Zmiz1^{fl/wt}$ (n=8) and $Zmiz1^{fl/wt;iECKO}$ (n=8) retinas at P7. Error bars represent mean ± s.e.m; two-tailed unpaired t-test. ns (not significant; P>0.05), *P<0.05, **P<0.01, ***P<0.001, ****P < 0.0001.

**Figure 3. Endothelial Zmiz1 is dispensable for vascular remodeling and maintenance during adulthood. A,** Schematic showing tamoxifen administration in 4 week old mice and analysis at 6 weeks. **B,** Representative images of IB4 immunolabeled whole-mount retinas at P42. Scale bars 1000 μm. **C,** Representative images of IB4 positive vessels in the superficial and deep layers of $Zmiz1^{fl/fl}$ and $Zmiz1^{iECKO}$ retinas at P42. Scale bars: 200 μm. **D** and **E,** Quantification of superficial and deep vascular plexus density in $Zmiz1^{fl/fl}$ (n=6) and $Zmiz1^{iECKO}$ (n=8) retinas at P42. Error bars represent mean ± s.e.m; two-tailed unpaired t-test. ns (not significant; P>0.05), *P<0.05, **P<0.01, ***P<0.001, ****P < 0.0001.

**Figure 4. Zmiz1 depletion in MS1 ECs leads to migration and sprouting defects. A,** Representative bright field images of control(non-silencing)-shRNA and *Zmiz1*-shRNA MS1 cells at 0 and 120 hours in the fibrin gel bead assay (red arrowheads mark vascular sprouts). Scale bars: 1 mm. **B,** Quantification of the number of sprouts per bead in control-shRNA (n=27) and *Zmiz1*-shRNA cells (n=30). **C,** Representative images of control-shRNA and *Zmiz1*-shRNA MS1 cells immunofluorescently stained for EdU (red) and DAPI (blue) following EdU assays. Scale bars: 100 μm. **D,** Quantification of percentage of EdU positive cells in both control-shRNA (n=3) and *Zmiz1*-shRNA (n=3) MS1 cells. **E,** Representative bright field images of control-shRNA and *Zmiz1*-shRNA MS1 cells at 0 and 16 hours post scratch (dashed red lines demarcate the EC and EC-free boundaries). Scale bars: 500 μm. **F,** Quantification of the percentage of wound closure in control-shRNA (n=3) and *Zmiz1*-shRNA (n=3) MS1 cells after 16 hours. **G,** *Zmiz1* mRNA levels in control-shRNA (n=3) versus *Zmiz1*-shRNA (n=3) cells via qRT-PCR. Error bars represent mean ± s.e.m; two-tailed unpaired t-test. ns (not significant; P>0.05), *P<0.05, **P<0.01, ***P<0.001, ****P < 0.0001.

**Figure 5. Loss of Zmiz1 in ECs results in transcriptomic downregulation of EC migration, angiogenesis and vascular development associated genes. A,** MA plot of differentially expressed genes obtained from RNA-seq data between control and *Zmiz1*-shRNA MS1 ECs. Red dots, upregulated genes (718); blue dots, downregulated genes (993). **B, C,** Top Gene ontology (GO) biological process terms enriched in upregulated genes (**B**) downregulated genes (**C**) (False discovery rate (FDR)≤ 0.05). **D,** Analysis of transcription factor (TF) regulated enriched genes within the downregulated genes data set (False discovery rate (FDR)≤ 0.05).

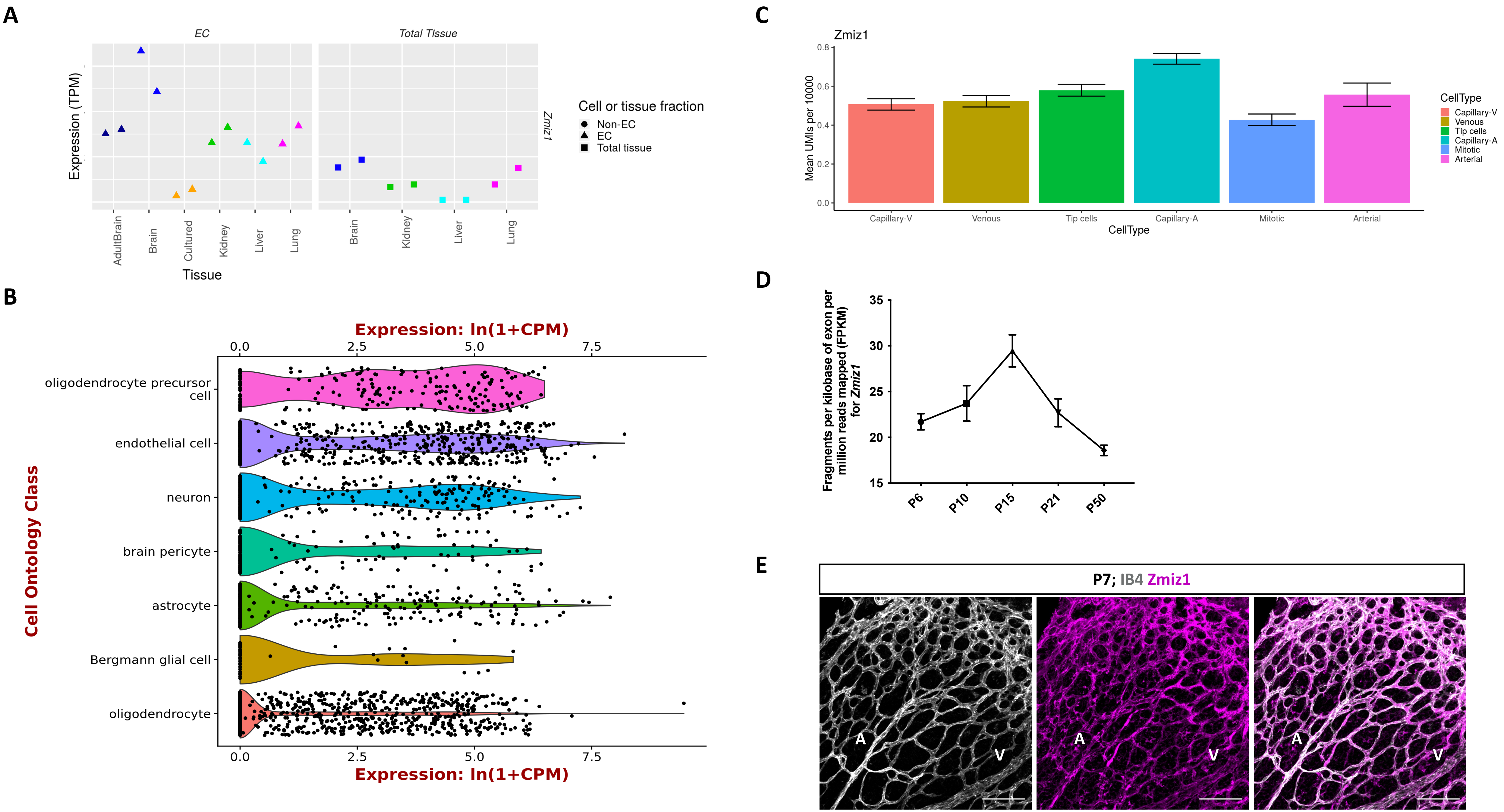

A
EC
Total Tissue
Expression (TPM)
Zmiz1
Cell or tissue fraction
Non-EC
EC
Total tissue
AdultBrain
Brain
Cultured
Kidney
Liver
Lung
Tissue
B
Expression: ln(1+CPM)
0.0
2.5
5.0
7.5
Cell Ontology Class
oligodendrocyte precursor cell
endothelial cell
neuron
brain pericyte
astrocyte
Bergmann glial cell
oligodendrocyte
C
Zmiz1
Mean UMIs per 10000
0.0
0.2
0.4
0.6
0.8
Capillary-V
Venous
Tip cells
Capillary-A
Mitotic
Arterial
CellType
D
Fragments per kilobase of exon per million reads mapped (FPKM) for Zmiz1
15
20
25
30
35
P6
P10
P15
P21
P50
E
P7; IB4 Zmiz1
A
V

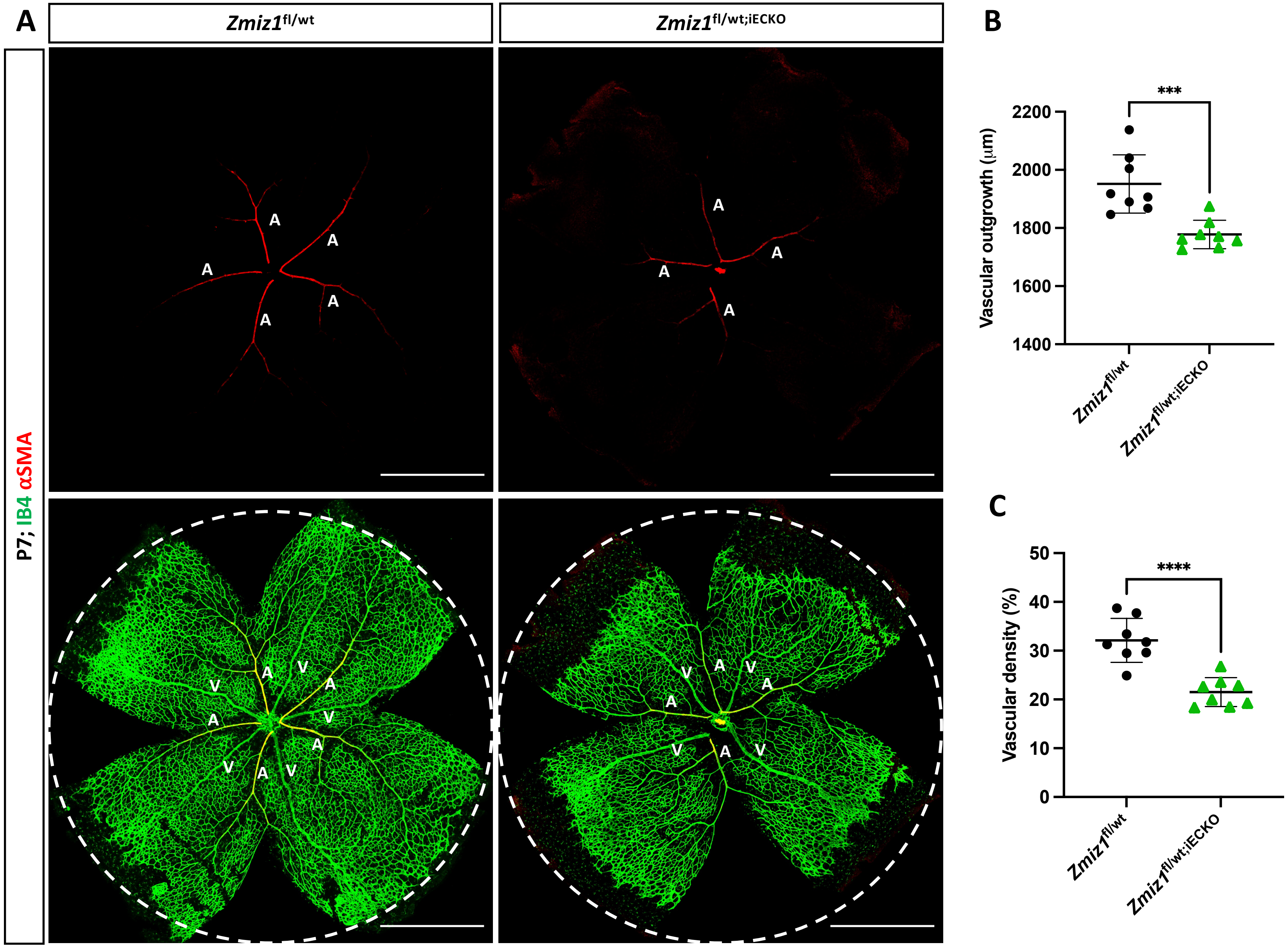

A
Zmiz1fl/wt
Zmiz1fl/wt;iECKO
P7; IB4 αSMA
A
V
B
***
Vascular outgrowth (μm)
2200
2000
1800
1600
1400
Zmiz1fl/wt
Zmiz1fl/wt;iECKO
C
****
Vascular density (%)
50
40
30
20
10
0
Zmiz1fl/wt
Zmiz1fl/wt;iECKO

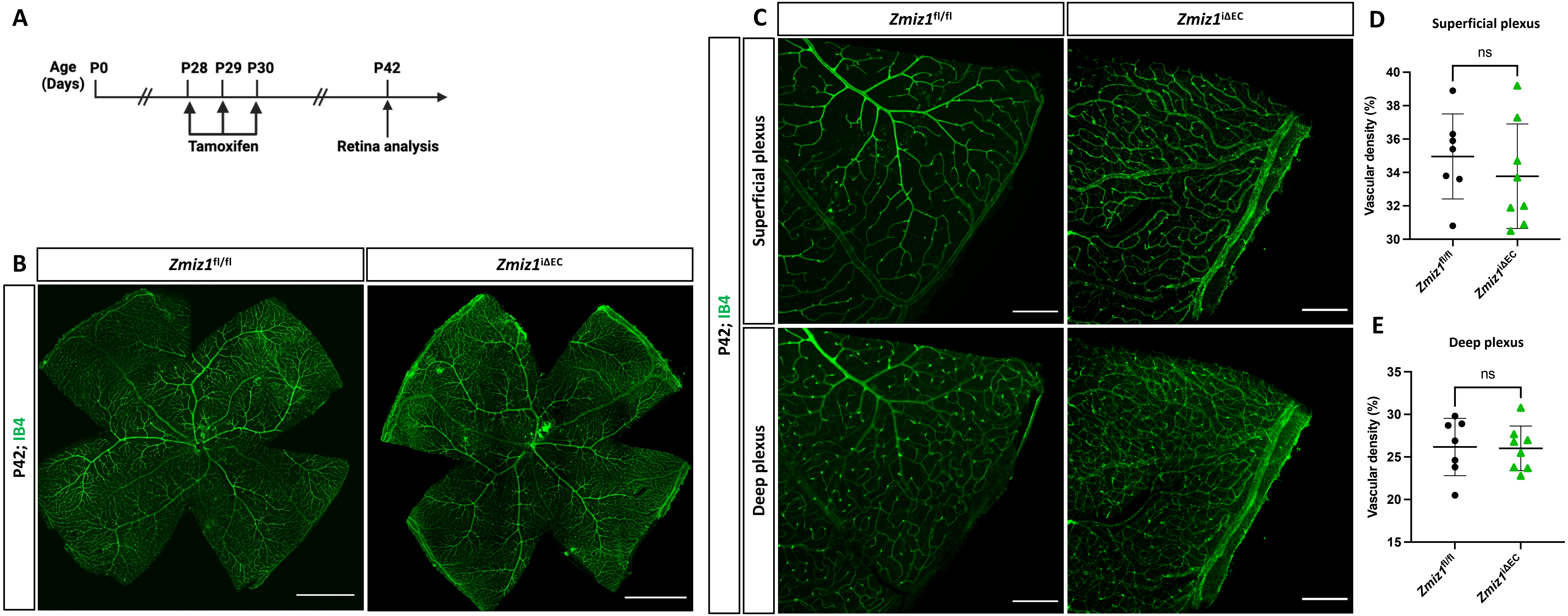

A
Age (Days)
P0
P28
P29
P30
P42
Tamoxifen
Retina analysis
B
Zmiz1fl/fl
Zmiz1iΔEC
P42; IB4
C
Superficial plexus
Deep plexus
D
Superficial plexus
ns
Vascular density (%)
40
38
36
34
32
30
E
Deep plexus
35
30
25
20
15

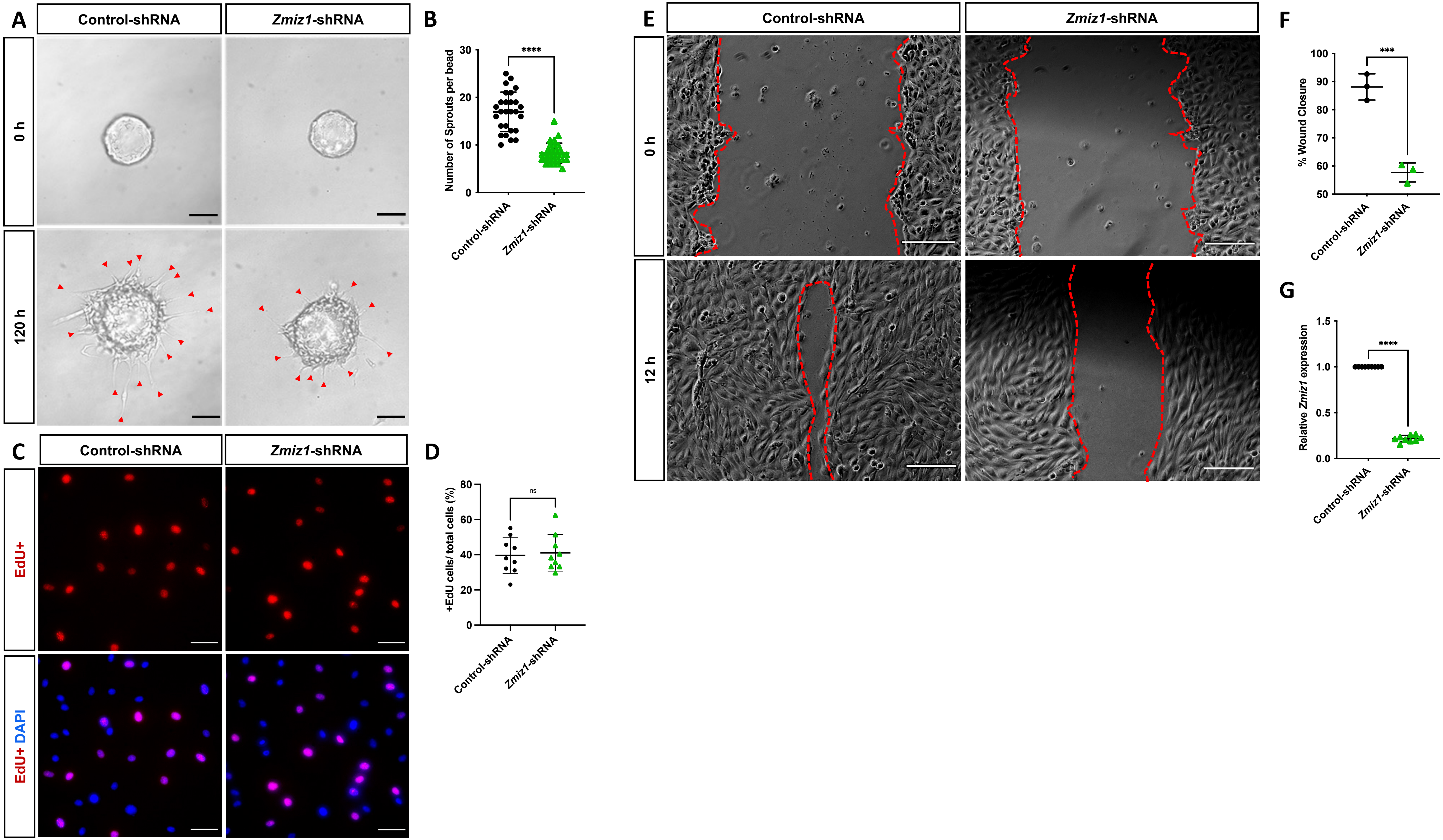
A
Control-shRNA
Zmiz1-shRNA
0 h
120 h
B
Number of Sprouts per bead
****
Control-shRNA
Zmiz1-shRNA
C
EdU+
EdU+ DAPI
D
+EdU cells/ total cells (%)
ns
E
12 h
F
% Wound Closure
***
G
Relative Zmiz1 expression
****

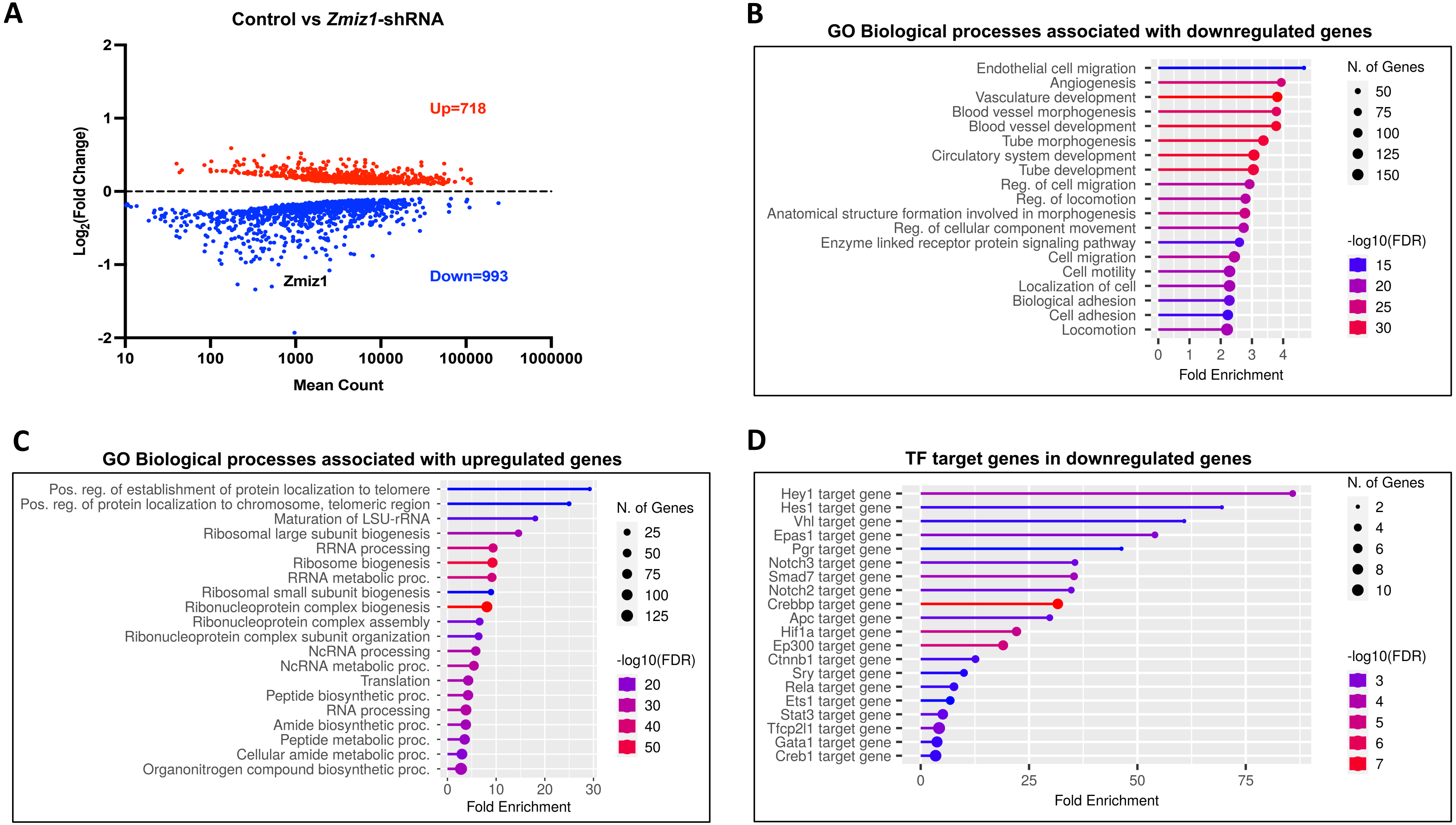
A
Control vs Zmiz1-shRNA
Up=718
Down=993
Zmiz1
Log2(Fold Change)
Mean Count
B
GO Biological processes associated with downregulated genes
Endothelial cell migration
Angiogenesis
Vasculature development
Blood vessel morphogenesis
Blood vessel development
Tube morphogenesis
Circulatory system development
Tube development
Reg. of cell migration
Reg. of locomotion
Anatomical structure formation involved in morphogenesis
Reg. of cellular component movement
Enzyme linked receptor protein signaling pathway
Cell migration
Cell motility
Localization of cell
Biological adhesion
Cell adhesion
Locomotion
Fold Enrichment
N. of Genes
-log10(FDR)
C
GO Biological processes associated with upregulated genes
Pos. reg. of establishment of protein localization to telomere
Pos. reg. of protein localization to chromosome, telomeric region
Maturation of LSU-rRNA
Ribosomal large subunit biogenesis
RRNA processing
Ribosome biogenesis
RRNA metabolic proc.
Ribosomal small subunit biogenesis
Ribonucleoprotein complex biogenesis
Ribonucleoprotein complex assembly
Ribonucleoprotein complex subunit organization
NcRNA processing
NcRNA metabolic proc.
Translation
Peptide biosynthetic proc.
RNA processing
Amide biosynthetic proc.
Peptide metabolic proc.
Cellular amide metabolic proc.
Organonitrogen compound biosynthetic proc.
Fold Enrichment
N. of Genes
-log10(FDR)
D
TF target genes in downregulated genes
Hey1 target gene
Hes1 target gene
Vhl target gene
Epas1 target gene
Pgr target gene
Notch3 target gene
Smad7 target gene
Notch2 target gene
Crebbp target gene
Apc target gene
Hif1a target gene
Ep300 target gene
Ctnnb1 target gene
Sry target gene
Rela target gene
Ets1 target gene
Stat3 target gene
Tfcp2l1 target gene
Gata1 target gene
Creb1 target gene
Fold Enrichment
N. of Genes
-log10(FDR)
